# Tissue-adjusted pathway analysis of cancer (TPAC)

**DOI:** 10.1101/2022.03.17.484779

**Authors:** H. Robert Frost

## Abstract

We describe a novel single sample pathway analysis method for cancer transcriptomics data named tissue-adjusted pathway analysis of cancer (TPAC). The TPAC method leverages information about the normal tissue-specificity of human genes to compute a robust multivariate distance score that quantifies pathway dysregulation in each profiled tumor. Because the null distribution of the TPAC scores has an accurate gamma approximation, both population and sample-level inference is supported. As we demonstrate through an analysis of gene expression data from The Cancer Genome Atlas (TCGA), TPAC pathway scores are more strongly associated with both patient prognosis and tumor stage than the scores generated by existing single sample pathway analysis methods. An R package implementing the TPAC method can be found at https://hrfrost.host.dartmouth.edu/TPAC.

## 1 Background

Cancer develops when somatic alterations disrupt pathways associated with genome maintenance, cell proliferation, or cell survival to give a relative growth advantage to the impacted cell population [1]. Given the central role of pathway dysregulation in tumor initiation, growth and metastatic spread, basic and translational cancer research is often focused on pathway-related questions, e.g., How do somatic alterations cause dysregulation of specific pathways? What is the impact of pathway dysregulation on patient prognosis? What types of therapeutic agents can restore normal pathway function? One common approach for answering these questions involves the analysis of tumor-derived genomic data using gene set testing, or pathway analysis, methods. Gene set testing is a hypothesis aggregation technique that evaluates statistics computed on functionally related groups of genes, e.g., the sets maintained in collections like the Molecular Signatures Database (MSigDB) [2]. By focusing on gene sets, rather than individual genes, pathway analysis can significantly improve power, interpretation and replication [3–6].

Although the pathways most commonly impacted in cancer have been identified [1] and progress has been made developing cancer-specific pathway analysis methods that can integrate gene expression and somatic alteration data [7–11], current approaches leverage just tumor-specific genomic data and do not take into account gene activity in the associated normal tissue. This is an important limitation given the strong association between gene activity in normal tissues and the pathophysiology of cancers originating in those tissues. Cancer biology is highly tissue and cell type-specific [12–16] with most driving somatic alterations either occuring in only a small number of cancer types, e.g., KRAS mutations in pancreatic, lung and colorectal cancers [17], or having a functional impact that varies between impacted tissues, e.g., germline BRCA1/BRCA2 mutations that only drive cancer in estrogen-sensitive tissues [18]. As we explored in a recent paper [19], this tissue-specificity means that the pattern of gene activity (as quantified by mRNA expression) in normal tissues carries important information regarding the biology of the associated cancer types. In this paper, we demonstrated through an analysis of solid tumor data from The Cancer Genome Atlas (TCGA) [20] and normal tissue data from the Human Protein Atla (HPA) [21] that the association between tissue-specific and cancer-specific expression values, i.e., the ratio of gene expression in a specific tissue or cancer relative to the mean across multiple tissues or cancers, can be used to improve survival analysis, the comparative analysis of distinct cancer types, and the analysis of cancer/normal tissue pairs. Specifically, we found that genes enriched in normal tissues are more likely to be down-regulated in the associated cancer with elevated expression associated with a favorable cancer prognosis (see the *ρ*_*cancer/norm*_ and *ρ*_*surv*_ columns in Table 1 for statistics that capture these associated for different cancer/normal pairs).

**Table 1:** The 21 TCGA cancer types and corresponding HPA normal tissues supported by TPAC. The *ρ*_*cancer/norm*_ column holds the Spearman rank correlation between normal tissue-specific gene weights (the log fold-change of the mean expression in the normal tissue to the mean in all normal tissues) and the log fold-change of the mean expression in the cancer type to mean expression in the normal tissue. The *ρ*_*surv*_ column holds the Spearman rank correlation between normal tissue-specific gene weights and the signed log of the p-value from a Kaplan-Meir test of the association between gene expression and cancer survival as computed by Uhlen et al. [21]), which is computed as -log(p-value) for favorable genes and log(p-value) for unfavorable genes. The contents of this table are a synthesis of information from Tables 2 and 3 in Frost [19].

| TCGA abbrev. | Cancer type | HPA tissue | $\rho_{cancer/norm}$ | $\rho_{surv}$ |
| --- | --- | --- | --- | --- |
| BLCA | Bladder Urothelial Carcinoma | urinary bladder | -0.299 | -0.0229 |
| BRCA | Breast Invasive Carcinoma | breast | -0.437 | -0.0993 |
| CESC | Cervical Squamous Cell Carcinoma | cervix, uterine | -0.432 | -0.105 |
| COAD | Colon Adenocarcinoma | colon | -0.057 | 0.304 |
| GBM | Glioblastoma Multiforme | cerebral cortex | -0.446 | 0.0764 |
| HNSC | Head and Neck Squamous Cell Carcinoma | tonsil | -0.48 | 0.11 |
| KICH | Kidney Chromophobe | kidney | -0.232 | 0.454 |
| KIRC | Kidney Renal Clear Cell Carcinoma | kidney | -0.436 | 0.454 |
| KIRP | Kidney Renal Papillary Cell Carcinoma | kidney | -0.311 | 0.454 |
| LIHC | Liver Hepatocellular Carcinoma | liver | -0.358 | 0.35 |
| LUAD | Lung Adenocarcinoma | lung | -0.363 | 0.073 |
| LUSC | Lung Squamous Cell Carcinoma | lung | -0.406 | 0.073 |
| OV | Ovarian Serous Cystadenocarcinoma | ovary | -0.546 | -0.0348 |
| PAAD | Pancreatic Adenocarcinoma | pancreas | -0.643 | 0.148 |
| PRAD | Prostate Adenocarcinoma | prostate | -0.179 | -0.0397 |
| READ | Rectum Adenocarcinoma | rectum | -0.208 | 0.349 |
| SKCM | Skin Cutaneous Melanoma | skin | -0.444 | -0.0707 |
| STAD | Stomach Adenocarcinoma | stomach | -0.258 | 0.344 |
| TGCT | Testicular Germ Cell Tumors | testis | -0.636 | 0.256 |
| THCA | Thyroid Carcinoma | thyroid gland | -0.441 | -0.0851 |
| UCEC | Uterine Corpus Endometrial Carcinoma | endometrium | -0.45 | 0.0276 |

To leverage these associations between normal tissue-specificity and cancer biology for pathway analysis, we have developed a new single sample pathway analysis method for tumor-derived transcriptomics data named TPAC (tissue-adjusted pathway analysis of cancer). The TPAC method leverages information about the normal tissue-specificity of human genes to compute a robust multivariate distance score that quantifies pathway dysregulation in each profiled tumor. TPAC currently supports the 21 solid tumor types listed in Table 1. Because the null distribution of the TPAC scores has an accurate gamma approximation, both population and sample-level inference is supported. As we demonstrate through an analysis of TCGA gene expression data, TPAC pathway scores are more strongly associated with both patient prognosis and tumor stage than the scores generated by existing single sample pathway analysis methods. In the remainder of this paper, we detail the TPAC method in Section 2, and characterize the performance of TPAC relative to existing single sample methods in Section 3. The companion website (https://hrfrost.host.dartmouth.edu/TPAC) provides access to related resources including an R package and vignette illustrating the application of TPAC to TCGA liver cancer RNA-seq data.

## 2 Methods

### 2.1 Data sources

The findings detailed in Section 3 are based on bulk RNA-seq and clinical data from The Cancer Genome Atlas (TCGA) [20] for 21 human solid cancers and bulk RNA-seq data from the Human Protein Atlas (HPA) [22] for the associated 18 normal human tissues. These cancer types and normal tissues are listed in Table 1. Alternative cancer outcomes (e.g., progression-free interval) were accessed from the TCGA Pan-Cancer Clinical Data Resource (TCGA-CDR) [23]. Gene set definitions were accessed from version 7.2 of the Molecular Signatures Database (MSigDB) [2]. Additional details on these data sources can be found in the SI Methods.

### 2.2 TPAC method

The TPAC method generates single sample gene set scores from tumor gene expression data using a modified version of the classic Mahalanobis multivariate distance measure [24]. TPAC takes four inputs:

- **X**: *n* × *p* matrix that holds the normalized expression measurements for *p* genes in *n* tumors all of the same type (e.g., normalized RNA-seq data from TCGA). For the current implementation of TPAC, the tumor type is limited to one of the 21 solid cancers listed in Table 1. Genes with 0 variance are removed.
- **t**: a length *p* vector that holds the mean gene expression values (as quantified by the HPA) for the normal tissue associated with the cancer type whose data is held in **X**.
- 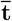: a length *p* vector that holds the average of the mean gene expression values across all 18 normal tissues in Table 1.
- **A**: *m* × *p* matrix that represents the annotation of *p* genes to *m* gene sets as defined by a collection from a repository like the Molecular Signatures Database (MSigDB) [2] (*a*_*i,j*_ = 1 if gene *j* belongs to gene set *i*).

Given **X, t**, 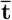, and **A**, TPAC computes *n* × *m* matrices **S, S**^+^, and **S**^−^. These matrices hold single sample scores for each of the *m* gene sets defined in **A** and *n* tumors captured in **X**. These single sample pathway scores are computed as follows:

1. **Compute normal tissue-specificity**: Let the length *p* vector **t**^*^ hold values representing the normal tissue-specificity of the *p* genes in **X**. These tissue-specific values are computed as 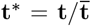, i.e., the fold-change in mean expression between the normal tissue associated with the target cancer type and the average in all 18 normal tissues listed in Table 1.
2. **Compute expression deviation between each tumor and associated normal tissue**: Let **Δ** hold the differences between the expression values in **X** and the normal tissue means in **t**, i.e, row *i* in **Δ** is computed by subtracting **t** from row *i* of **X**. Define two versions of **Δ, Δ**^+^ and **Δ**^−^, that capture just the positive or just the negative deviations. Specifically, element 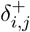 of **Δ**^+^ is set to element 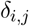 of **Δ** if *δ*_*i,j*_ ≥ 0, otherwise, it is set to 0. Similarily, element 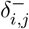 of **Δ**^−^ is set to element *δ*_*i,j*_ of **Δ** if *δ*_*i,j*_ *<* 0, otherwise, it is set to 0.
3. **Compute weighted sample covariance matrices**: Two weighted versions of the unbiased sample covariance matrix, 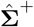 and 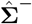, are computed that adjust the sample variance according to normal tissue-specificity. Specifically, diagonal element 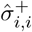 of 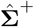 is set to the sample variance of gene *i* as computed on **X** multiplied by 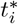 (the tissue-specificity value for gene *i*). Similarly, diagonal element 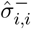 of 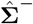 is set to the sample variance of gene *i* divided by 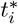. All off-diagonal elements in 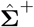 and 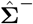 are set to 0. Table 2 captures the impact of this weighting on the sample variance for gene *i* (i.e., 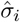), e.g., the variance is inflated for genes that are up-regulated in the normal tissue relative to other tissues (i.e., 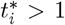) and have elevated expression in the tumor relative to normal tissue (i.e., *δ*_*i,j*_ ≥ 0). The impact of these variance changes is discussed in more detail in Section 2.3 below.

**Table 2:** Impact of tissue-specific weighting on sample variance for gene *i*.
4. **Compute modified Mahalanobis distances for positive expression deviations**: Let **M**^+^ be an *n* × *m* matrix of squared values of modified Mahalanobis distances. Each column *k* of **M**^+^, which holds the positive component of the sample-specific squared distances for gene set *k*, is calculated as:

| | $t_i^* \geq 1$ | $t_i^* < 1$ |
| --- | --- | --- |
| $\delta_{i,j} \geq 0$ | $\hat{\sigma}_i \uparrow$ | $\hat{\sigma}_i \downarrow$ |
| $\delta_{i,j} < 0$ | $\hat{\sigma}_i \downarrow$ | $\hat{\sigma}_i \uparrow$ |

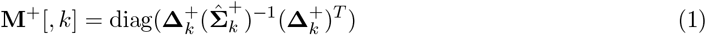

where *g* is the size of gene set *k*, 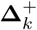 is a *n* × *g* matrix containing the *g* columns of **Δ**^+^ corresponding to the members of set *k*, and 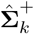 is a *g* × *g* matrix containing the *g* rows and columns of 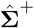 corresponding to the members of set *k*.
5. **Compute modified Mahalanobis distances for negative expression deviations**: Let **M**^−^ be an *n* × *m* matrix of squared values of modified Mahalanobis distances. Similar to **M**^+^, each column *k* of **M**^−^, which holds the negative component of the sample-specific squared distances for gene set *k*, is calculated as:

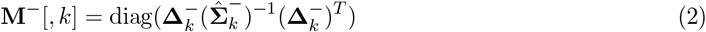

where *g* is the size of gene set *k*, 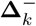 is a *n* × *g* matrix containing the *g* columns of **Δ**^−^ corresponding to the members of set *k*, and 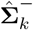 is a *g* × *g* matrix containing the *g* rows and columns of 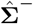 corresponding to the members of set *k*.
6. **Compute modified Mahalanobis distances for positive and negative expression deviations**: Let **M** be an *n* × *m* matrix of squared values of modified Mahalanobis distances that capture both positive and negative expression deviations from the associated normal tissue. These total squared distances are simply computed as the sum of the positive and negative distances:

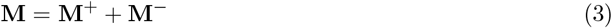
7. **Compute modified Mahalanobis distances on permuted X**: To capture the distribution of the squared modified Mahalanobis distances under the *H*_0_ that the expression values in **X** are uncorrelated with no mean difference between samples, the **M, M**^+^, and **M**^−^ matrices are recomputed on a version of **X** where the row labels of each column are randomly permuted. Let **X**_**p**_ represent the row-permuted version of **X** and let **M**_*p*_, 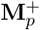, and 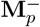 that hold the squared modified Mahalanobis distances computed on **X**_**p**_ according to (1), (2), or (3).
8. **Fit gamma distribution to columns of M**_*p*_, 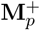, and 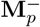: A separate gamma distribution is fit using the method of maximum likelihood (as implemented by the *fitdistr()* function in the MASS R package [25]) to the elements in each column of **M**_*p*_, 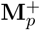, and 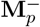. Let 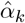 and 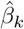, *k* ∈ 1, …, *m* represent the gamma shape and rate parameters estimated for gene set *k* on **M**_*p*_ using this procedure. For 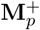 and 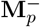, these estimated parameters are 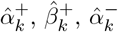, and 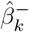.
9. **Use gamma cumulative distribution function (CDF) to compute tumor-specific gene set scores**: The tumor-specific gene set scores are set to the gamma CDF value for each element of **M, M**^+^, and **M**^−^. Specifically, each column *k* of **S**, which holds the tumor-specific scores for gene set *k*, is calculated as:

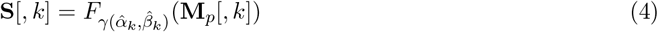

where 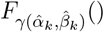 is the CDF for the gamma distribution with shape 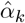 and rate 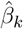. Under the *H*_0_ of uncorrelated expression, valid p-values can be generated by subtracting the elements of **S** from 1. The elements of **S**^+^ and **S**^−^ are populated using the same approach for the elements of **M**^+^ and **M**^−^ and gamma distributions fit on 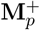, and 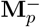.

The TPAC method is motivated in part by our previously developed Variance-adjusted Mahalanobis (VAM) method [26], which uses a modified Mahalanobis distance for cell-level gene set testing of single cell RNA-sequencing data. For the VAM approach, only positive distances, measured relative to the origin, are used and the sample covariance matrix is modified to capture just the technical component of gene expression variance. Similar to the use of gamma CDF values for the VAM method, the use of 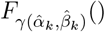 to generate TPAC scores has several important benefits: 1) it enables gene set inference for individual tumors, 2) it transforms the distances for gene sets of different sizes into a common scale, and 3) it produces scores that are bound between 0 and 1 and robust to large expression values.

### 2.3 Choice of S, S^+^ or S^−^

The three different TPAC generated score matrices, **S, S**^+^ and **S**^−^, capture distinct features of pathway dysregulation within each tumor and the choice of which scores to use will therefore vary depending on the analysis goals. For all matrices, large scores correspond to tumors that are more significantly dysregulated relative to the corresponding normal tissue and are therefore more likely on average to be associated with a poor prognosis or advanced cancer stage.

- **S**^+^: Large values in **S**^+^ correspond to tumors where expression of pathway genes is elevated relative to the associated normal tissue. The use of normal tissue-specificity to adjust sample variances (as detailed in Table 2 above) will prioritize expression differences for genes that are normally surpressed in the associated normal tissue, i.e., a tumor is considered more dysregulated if genes that are expressed at a low level in the associated normal tissue relative to other tissues are up-regulated in the tumor.
- **S**^−^: Large values in **S**^−^ correspond to tumors where expression of pathway genes is down-regulated relative to the associated normal tissue. In constrast to the impact on **S**^+^, the use of normal tissuespecificity to adjust sample variances leads to larger **S**^−^ values when tissue-specific genes are downregulated in the tumor, i.e., a tumor is considered more dysregulated if genes that are expressed at a high level in the associated normal tissue relative to other tissues are down-regulated in the tumor.
- **S**: Large values in **S** correspond to tumors where expression of pathway genes exhibit a combination of up and down-regulation relative to the associated normal tissue.

For most of the analysis results presented below, we use the scores in the **S** matrixsince these capture a wider range of dysregulation patterns.

### 2.4 Comparison methods

We compared the performance of the TPAC method against three existing single sample gene set testing techniques: GSVA [27], ssGSEA [28] and the z-scoring method of Lee et al. [29]. For all three methods, the implementation in the GSVA R package was used with default parameter settings. We also included results for a “null” version of TPAC as a negative control (i.e., TPAC scores with permuted sample labels to break any associations with cancer prognosis or stage), and a version of TPAC that does not use tissue-specific weights to adjust gene expression sample variances.

### 2.5 TCGA analyses

The TPAC method and the comparative methods outlined above in Section 2.4 were used to generate single sample gene set scores for TCGA RNA-seq data from the 21 cohorts listed in Table 1 and the 50 gene sets from the MSigDB Hallmark collection. The TPAC **S** matrix scores and scores from the comparative methods were used for the following analyses:

- **Landscape of pan-cancer pathway dysregulation**: Single sample pathway scores for tumors from all TCGA cohorts and Hallmark pathways were clustered and visualized to explore the pattern of pathway dysregulation across multiple cancer types.
- **Survival analysis**: Univariable Cox proportion hazards models were fit for each cohort using progression free interval (PFI) as the outcome and single sample pathway scores as a single predictor variable.
- **Tumor/lymph node stage analysis**: For each TCGA cohort, a Wilcoxon rank sum test was performed on the single sample scores generated by each method for all Hallmark pathways. For the analysis of tumor stage, the TCGA tumor stage was discretized as T01 vs non-T01. For the analysis of lymph node stage, the TCGA lymph node stage was discretized as N0 vs non-N0.
- **Single tumor inference**: To evaluate the use of TPAC scores for tumor-level inference, the CDF values in the **S** matrices for all cohorts were converted to p-values. False discovery rate (FDR) values were then computed for the family containing all pathway scores across the 21 TCGA cohorts and 50 Hallmark pathways using the method of Benjamini and Hochberg method [30].
- **Kaplan-Meir analysis**: A Kaplan-Meir analysis was performed for the TCGA KIRP cohort and the progression-free interval (PFI) outcome with patients stratified according to the significance of TPAC score for the MSigDB Hallmark MYC Targets V1 pathway. Significance was determined according to whether the FDR value associated with the TPAC score was *<* 0.25 where the family of hypotheses included the TPAC scores for all 50 Hallmark pathways for all 321 KIRP samples with PFI data (16,050 total hypotheses).

## 3 Results and Discussion

### 3.1 Type I error control and power

To explore the statistical properties of the TPAC method, multivariate normal data was simulated under both the null and alternative hypotheses. To estimate the type I error rate, we generated 10,0000 independent samples from a 40-dimensional multivate normal distribution with ***μ*** = **0** and **∑** = **I**. For this null simulation, p-values derived from the **S** matrix generated by TPAC for a single set containing all 40 variables maintained a type I error rate of 0.0498 at *α* = 0.05. To estimate power, a similar 10,000 by 40 MVN matrix was simulated with a constant offset of *δ* ∈ *{*0.1, 0.2, …, 2*}* added to the values for the first 1,000 samples. Figure 1 illustrates the estimated empirical power for different effect sizes at *α* = 0.05.

**Figure 1:**
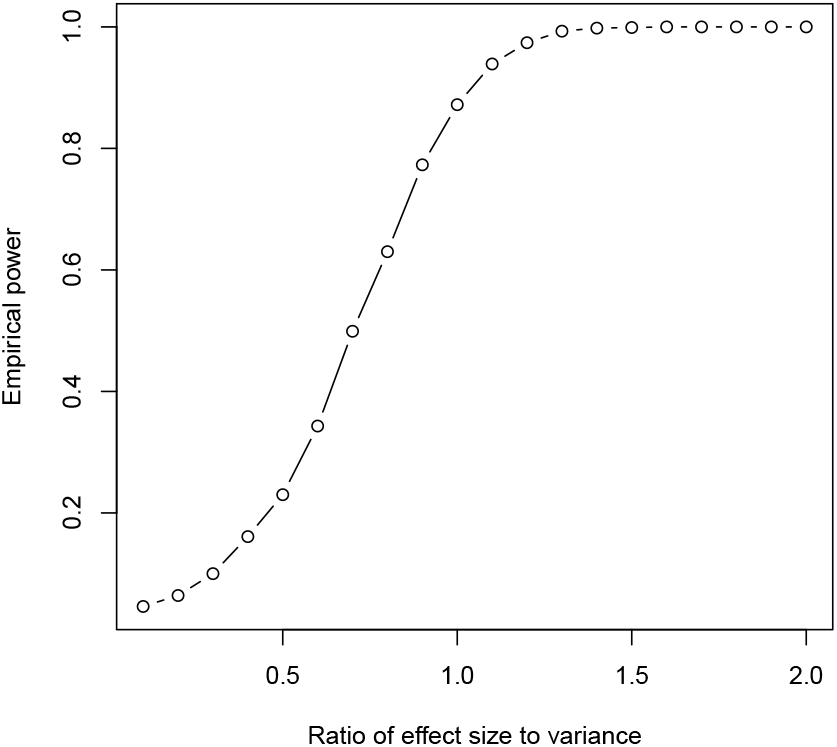
TPAC empirical power for different simulated effect sizes.

### 3.2 Pan-cancer distribution of TPAC scores

Figure 2 illustrates the overall pattern of **S** matrix TPAC scores for the MSigDB Hallmark pathways across tumors from all 21 evaluated TCGA cohorts. As seen in this figure, tumors cluster into four primary groups according to TPAC scores:

**Figure 2:**
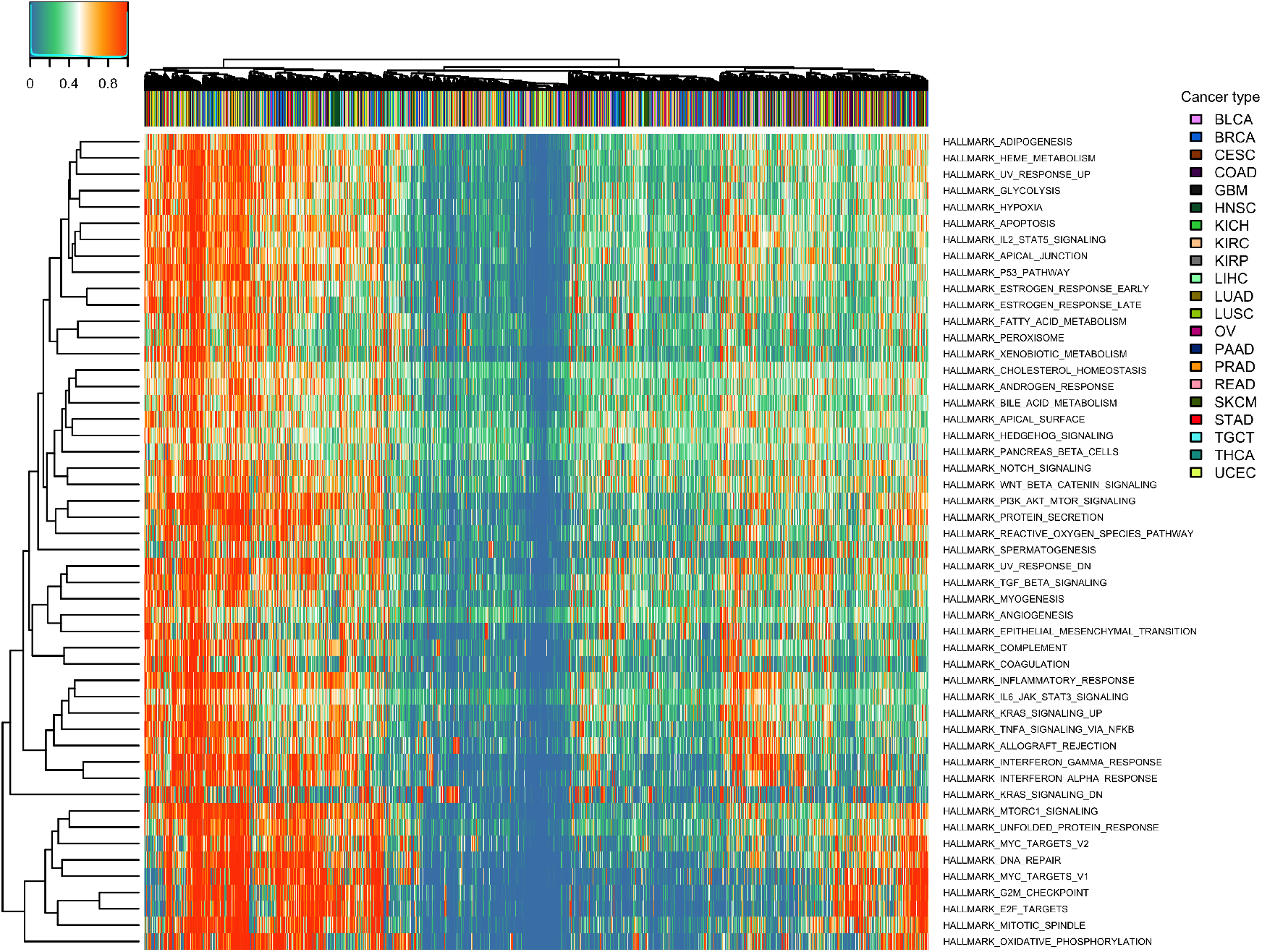
Heatmap illustrating the pan-cancer distribution of **S** matrix TPAC scores for the MSigDB Hallmark pathways.

1. Tumors that are highly dysregulated across all Hallmark pathways (i.e., tumors represented by the left-most columns in the heatmap)
2. Tumors that show pronounced dysregulation among proliferation pathways (i.e., tumors represented by the right-most columns in the heatmap)
3. Tumors that show pronounced dysregulation among pathways related to immune cell signaling (i.e., tumors represented by columns to the immediate left of the proliferation dysregulated tumors)
4. Tumors that exhibit limited gene expression dysregulation (i.e., tumors represented by columns in the middle of the heatmap).

### 3.3 Association between TPAC scores and cancer prognosis

Figure 3 illustrates the distribution of p-values from univariable Cox proportional hazards models fit for each TCGA cohort that use progression-free interval (PFI) as the outcome. Separate Cox models were fit for each combination of TCGA cohort, Hallmark pathway, and pathway analysis method with the distribution of all p-values associated with each analysis methods plotted as a separate curve. As shown in the figure, models using TPAC-generated pathway predictors provided the most significant associations, as determined by Cox models with FDR values ≤ 0.1. The magnitude and direction of the PFI associations for the TPAC-generated pathway scores are visualized in Figure 4. As illustrated by Figure 4, pathway dysregulation is usually associated with an unfavorable cancer prognosis and the strength of the association varies across the TCGA cohorts.

**Figure 3:**
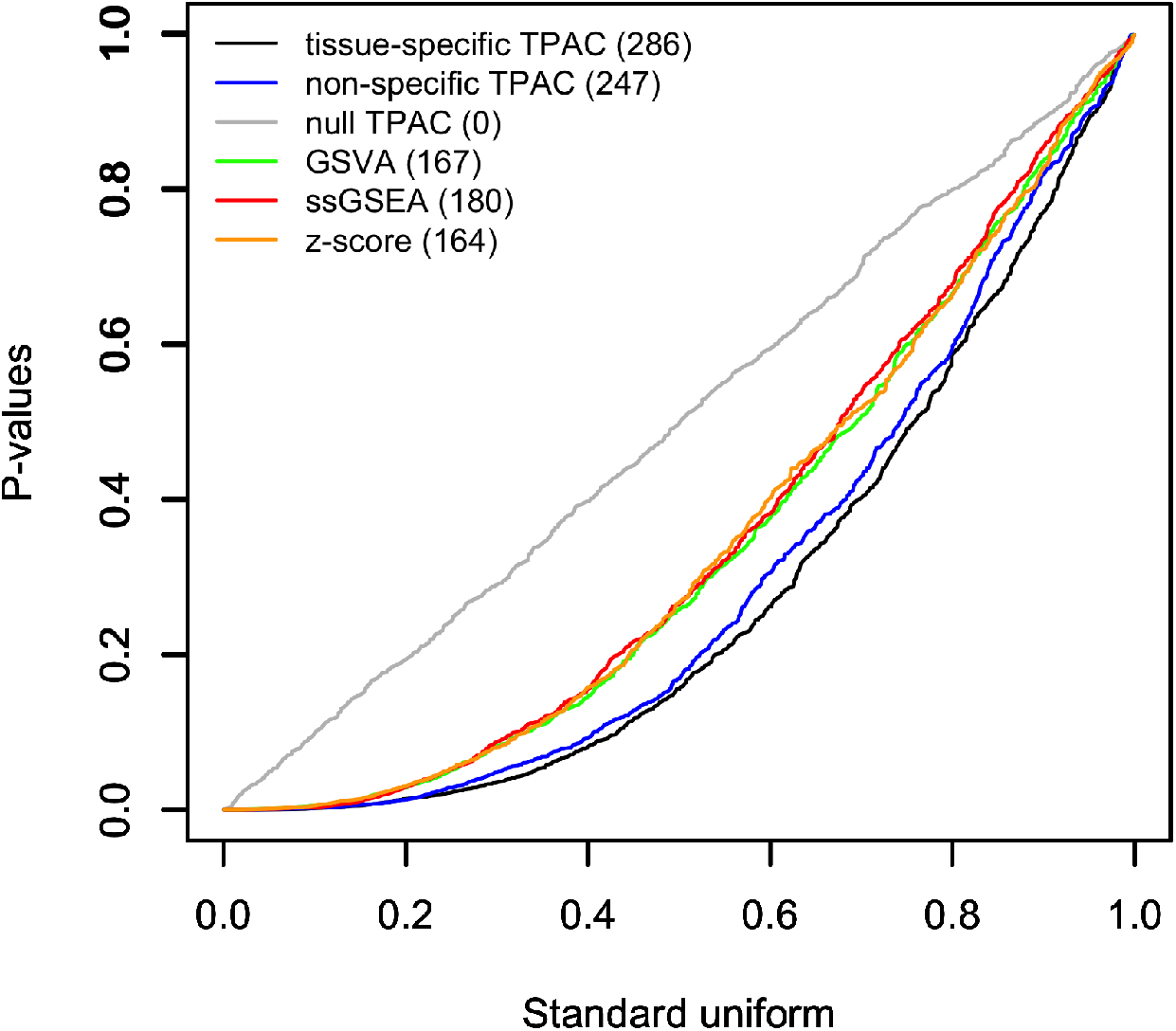
Q-Q plot comparing p-values from Cox proportional hazards models that use single sample pathway scores as the single predictor and PFI as the outcome against the *U* (0, 1) distribution expected under the null. The results for each evaluated single sample pathway analysis methods are plotted separately with the number of hypothesis tests associated with FDR values ≤ 0.1 listed in paratheses after the method in the legend.

**Figure 4:**
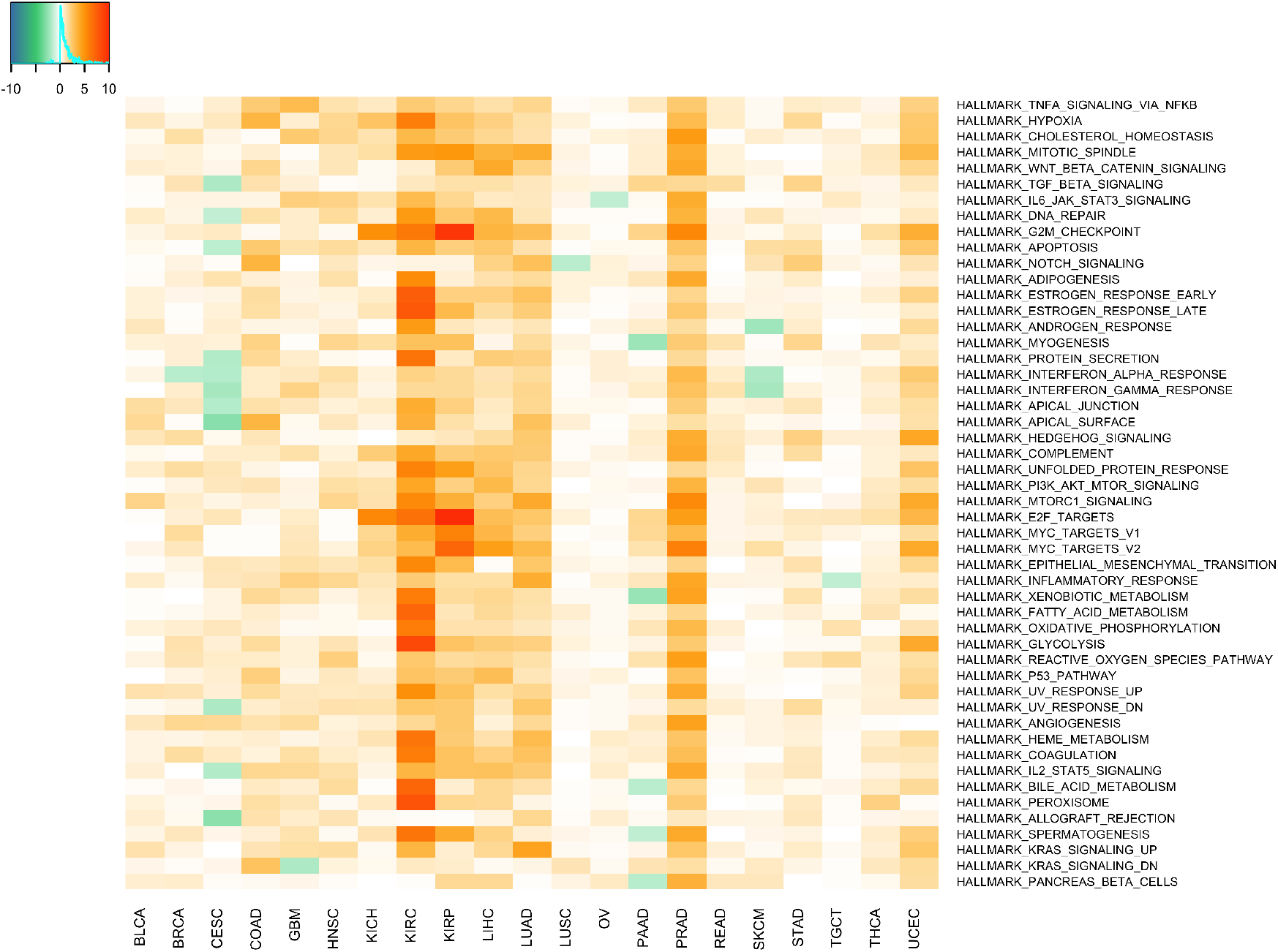
Visualization of p-values and association direction for univariable Cox proportional hazards models fit using TPAC-generated scores for Hallmark pathways as single predictors and PFI as the outcome. Each cell is colored according to the magnitude of the -log(p-value) from the Cox model with positive values for hazard ratios *>*= 1 and negative values for hazard rations *<* 1.

### 3.4 Association between TPAC scores and tumor stage

Figures 5 and 6 provide a similar visualization as Figures 3 and 4 of the p-values from statistical models based on single sample pathway scores computed using TPAC and the other comparison methods. For these figures, the p-values capture the association between pathway scores and discretized tumor stage (T01 vs. other) as evaluated using a Wilcoxon rank sum test. Although both TPAC variations yield more significant results than the comparison single sample methods, the version of TPAC that does not include an adjustment for tissue-specificity outperformed the version that includes the tissue-specific adjustment.

**Figure 5:**
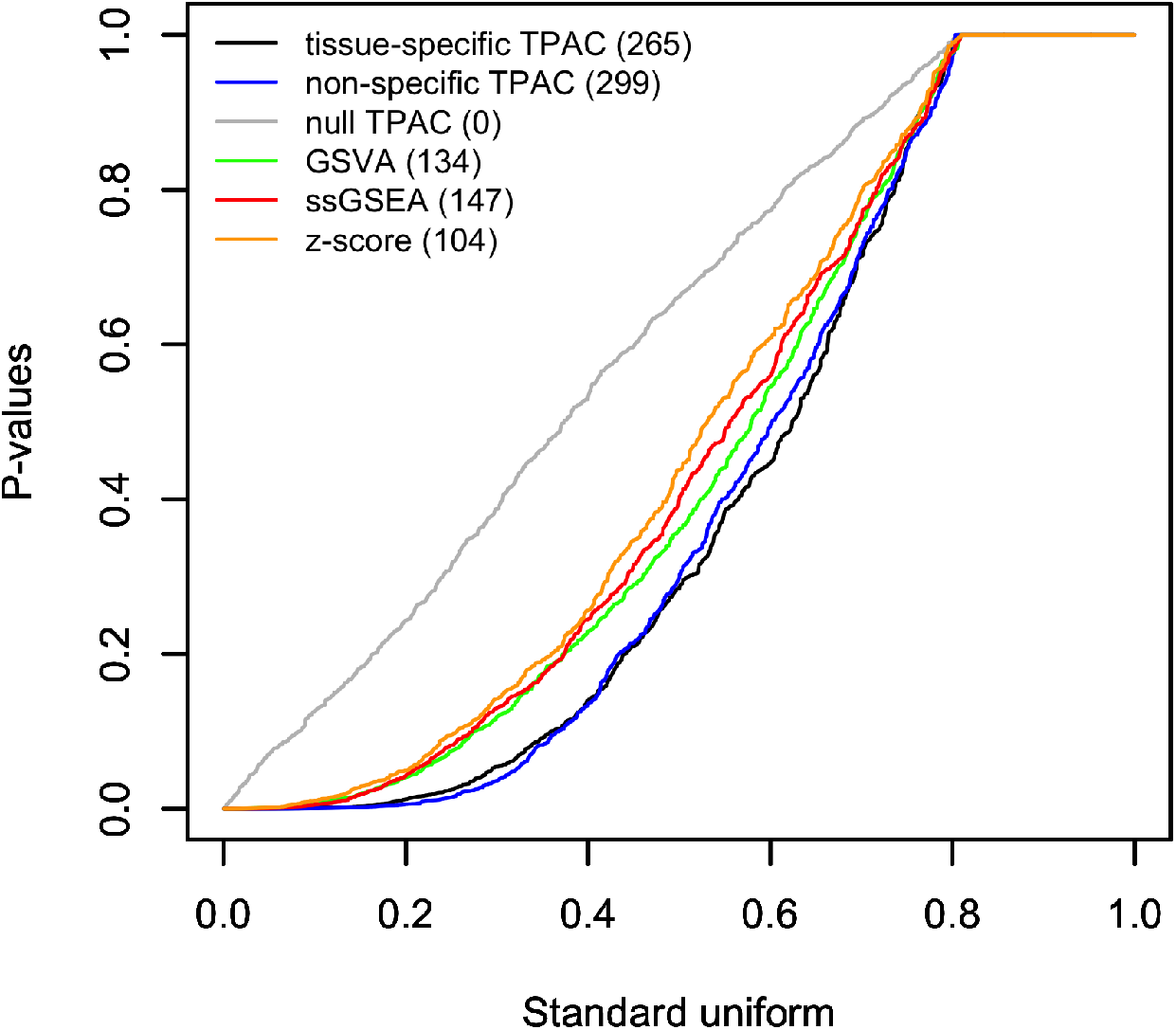
Q-Q plot comparing the distribution of p-values from Wilcoxon rank sum tests comparing single sample pathway scores for tumors with stage T01 vs. the scores for tumors with higher stages. The results for each evaluated single sample pathway analysis methods are plotted separately with the number of hypothesis tests associated with FDR values ≤ 0.1 listed in paratheses after the method in the legend.

**Figure 6:**
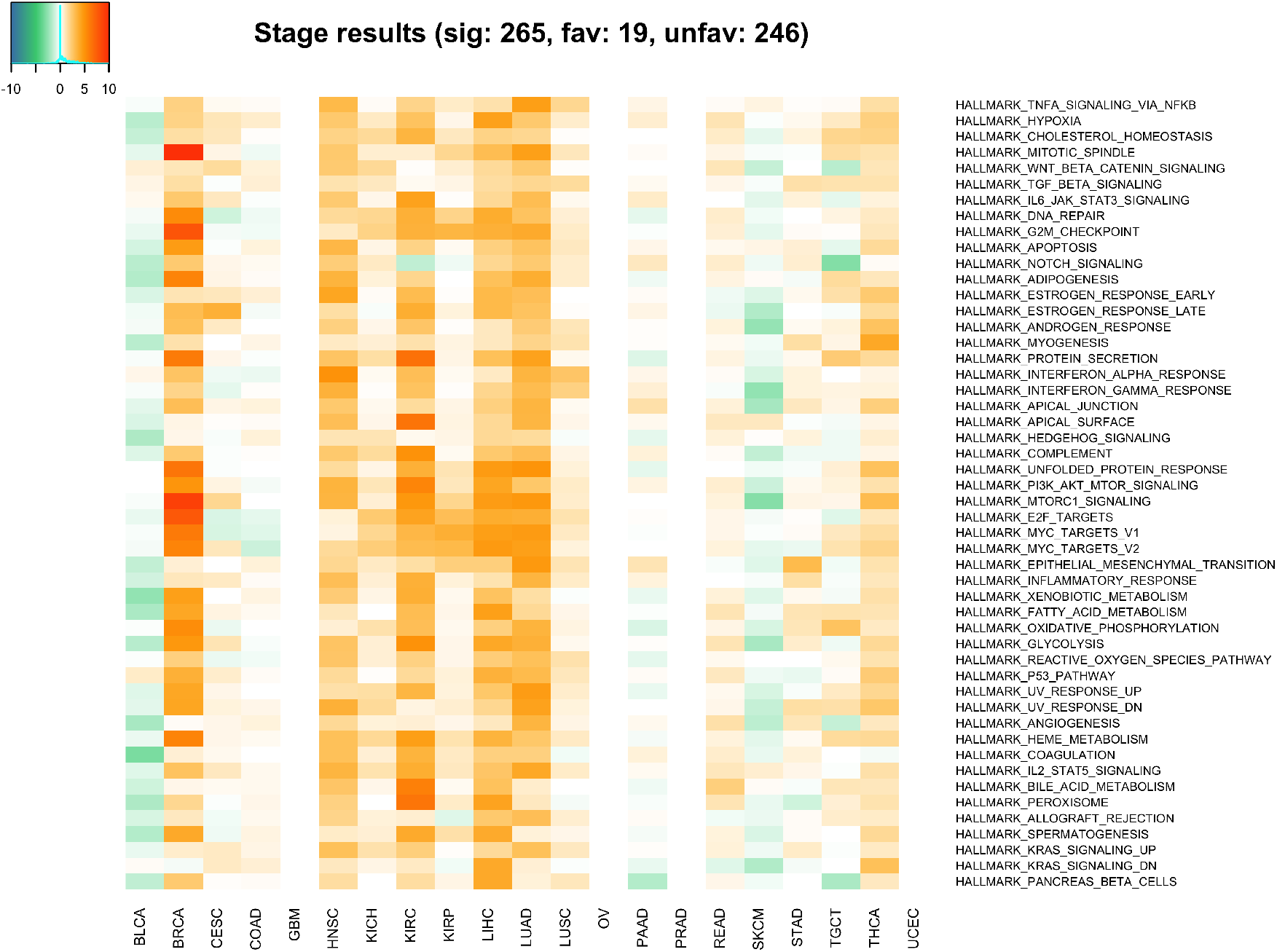
Visualization of p-values and association direction for Wilcoxon rank sum tests comparing TPAC scores for tumors with stage T01 vs. the scores for tumors with higher stages. Each cell is colored according to the magnitude of the -log(p-value) from the with positive values for cases where larger TPAC scores are associated with more advanced tumor stages and negative values where larger TPAC scores are associated with less severe tumor stages.

### 3.5 Association between TPAC scores and lymph node stage

Similar to Figures 5 and 6, Figures 7 and 8 visualize the association between single sample pathway scores and lymph node stage associated with each tumor. For the lymph node stage analysis, the standard TPAC generated substantially more significant associations (173) than any of the other comparison methods (from 108 to 118 findings).

**Figure 7:**
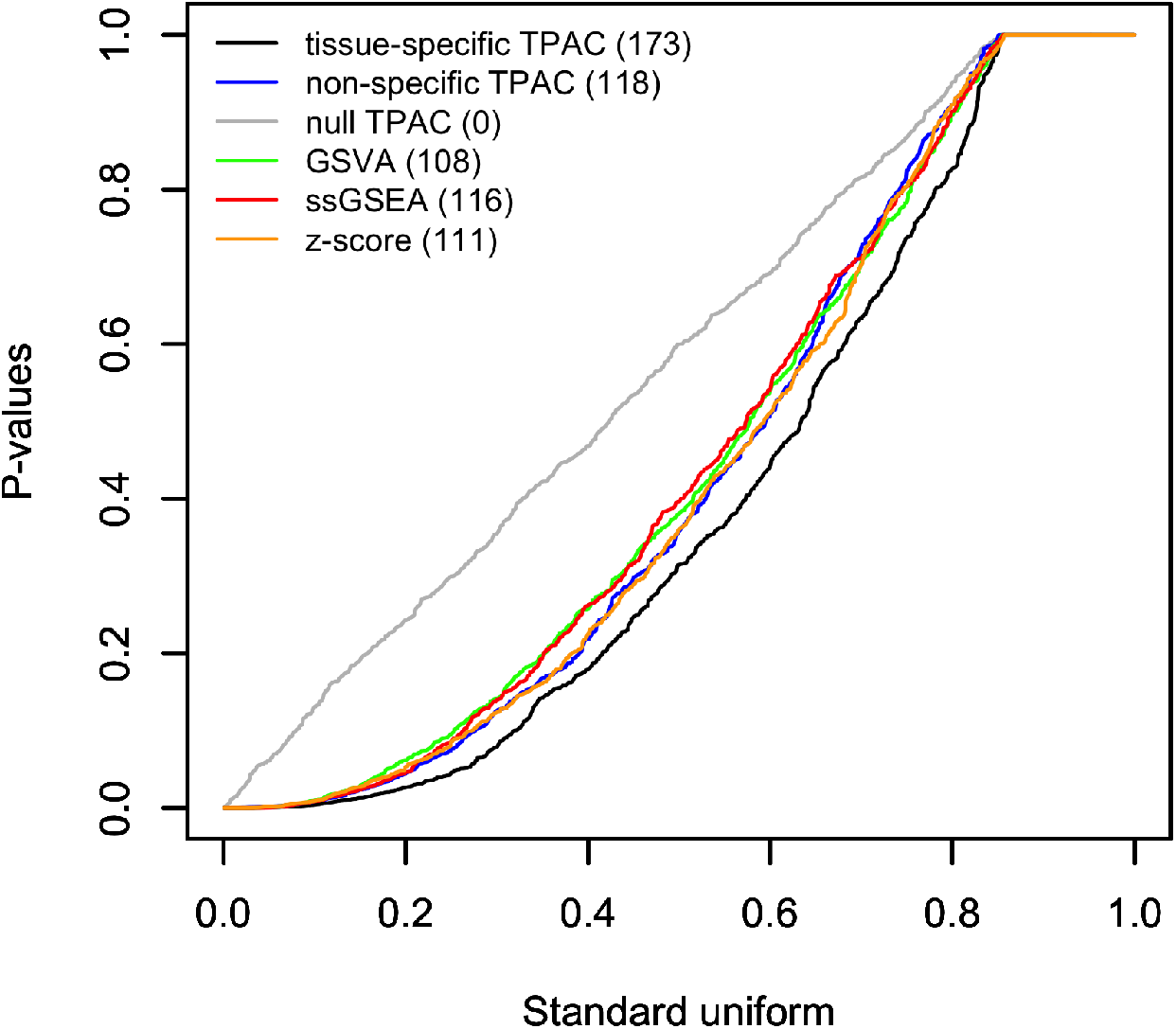
Q-Q plot comparing the distribution of p-values from Wilcoxon rank sum tests comparing single sample pathway scores for tumors associated with lymph node stage N0 vs. the scores for tumors associated with higher lymph node stages. The results for each evaluated single sample pathway analysis methods are plotted separately with the number of hypothesis tests associated with FDR values ≤ 0.1 listed in paratheses after the method in the legend.

**Figure 8:**
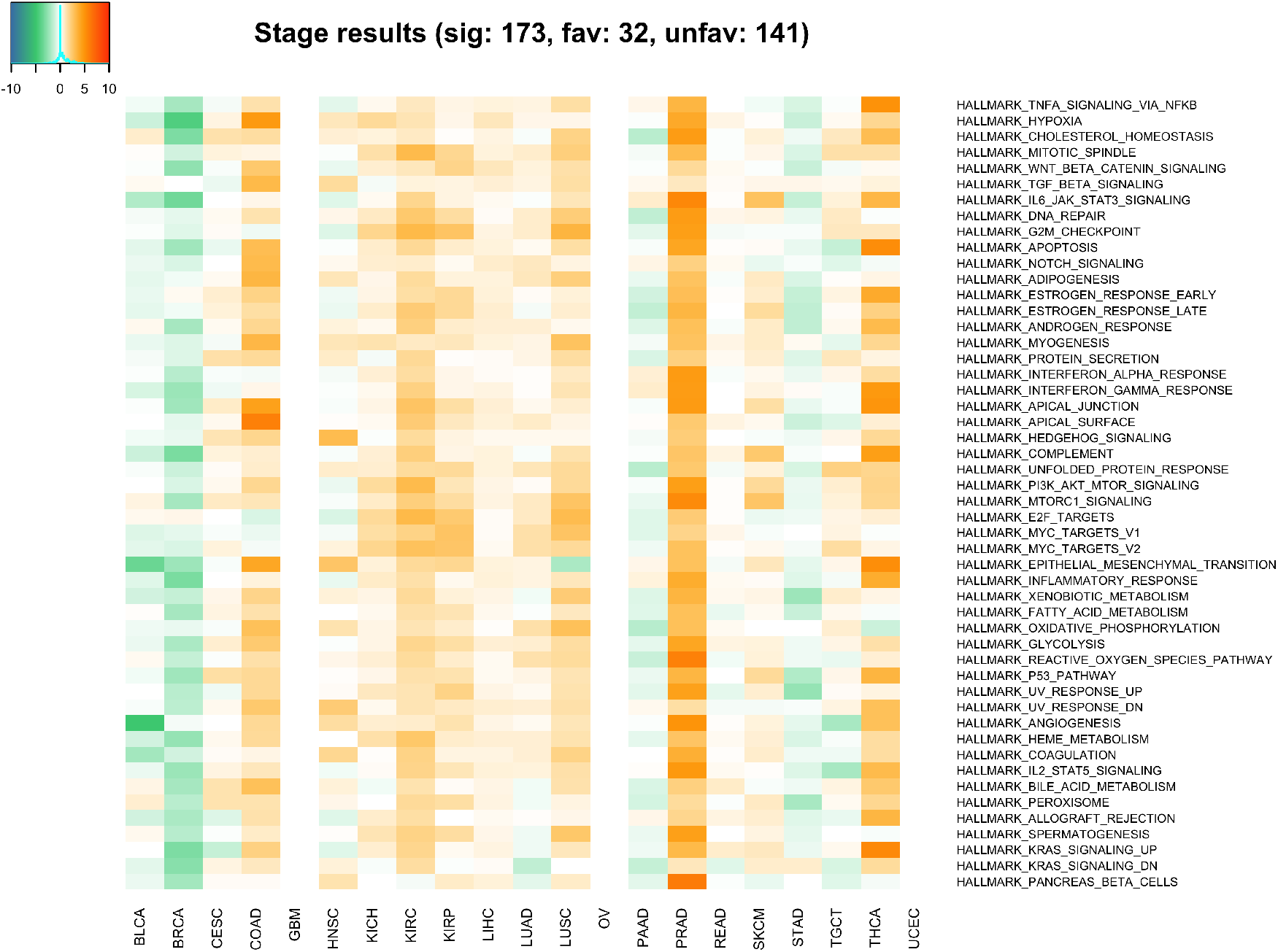
Visualization of p-values and association direction for Wilcoxon rank sum tests comparing TPAC scores for tumors associated with lymph node stage N0 vs. the scores for tumors associated with higher lymph node stages. Each cell is colored according to the magnitude of the -log(p-value) from the with positive values for cases where larger TPAC scores are associated with more advanced lymph node stages and negative values where larger TPAC scores are associated with less severe lymph node stages.

### 3.6 Sample-level inference

Figure 9 illustrates the use of TPAC scores for tumor-specific inference regarding pathway dysregulation. To generate this heatmap, the CDF values in the **S** matrices for all TCGA cohorts were converted to p-values. False discovery rate (FDR) values were then computed for the family containing all pathway scores across the 21 TCGA cohorts and 50 Hallmark pathways using the method of Benjamini and Hochberg method [30].

**Figure 9:**
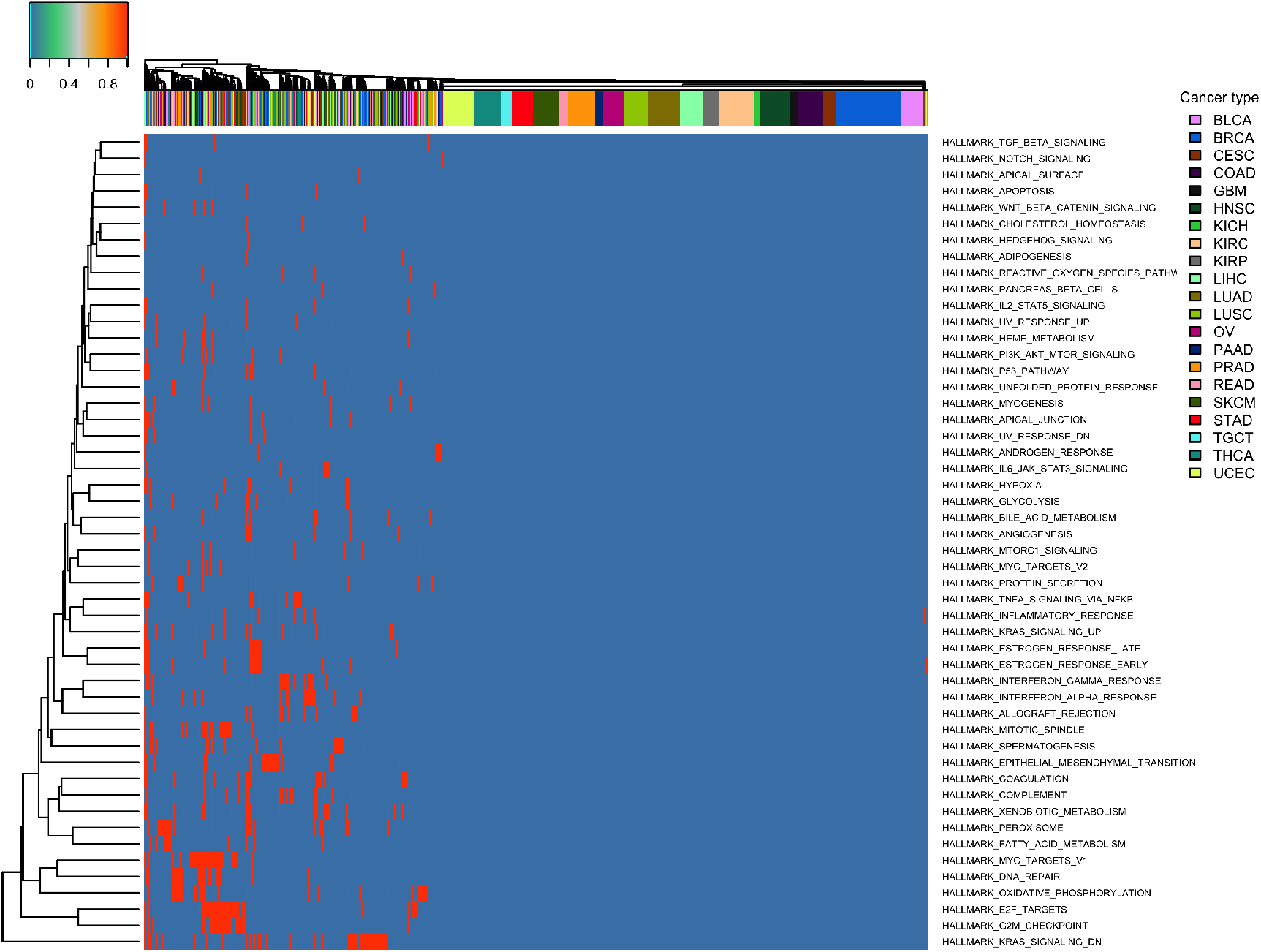
Illustration of TPAC score significance. If the FDR value for the TPAC score associated with a given tumor and pathway is ≥ 0.3, the value score is set to 0.

For cells corresponding to FDR values ≥ 0.3, the TPAC scores are set to 0 and the modified TPAC scores are then visualized as a heatmap. Figure 10 visualizes the use of tumor-level inference for survival analysis. Specifically, the TPAC scores for the Hallmark MYC Targets V1 pathway were discretized according to an FDR threshold of 0.25 and these discretized values were then used for a Kaplan-Meir analysis relative to patient PFI. As shown in this plot, tumors with significant dysregulation of the MYC Targets V1 pathway have a significantly worse prognosis than tumors lacking significant dysregulation.

**Figure 10:**
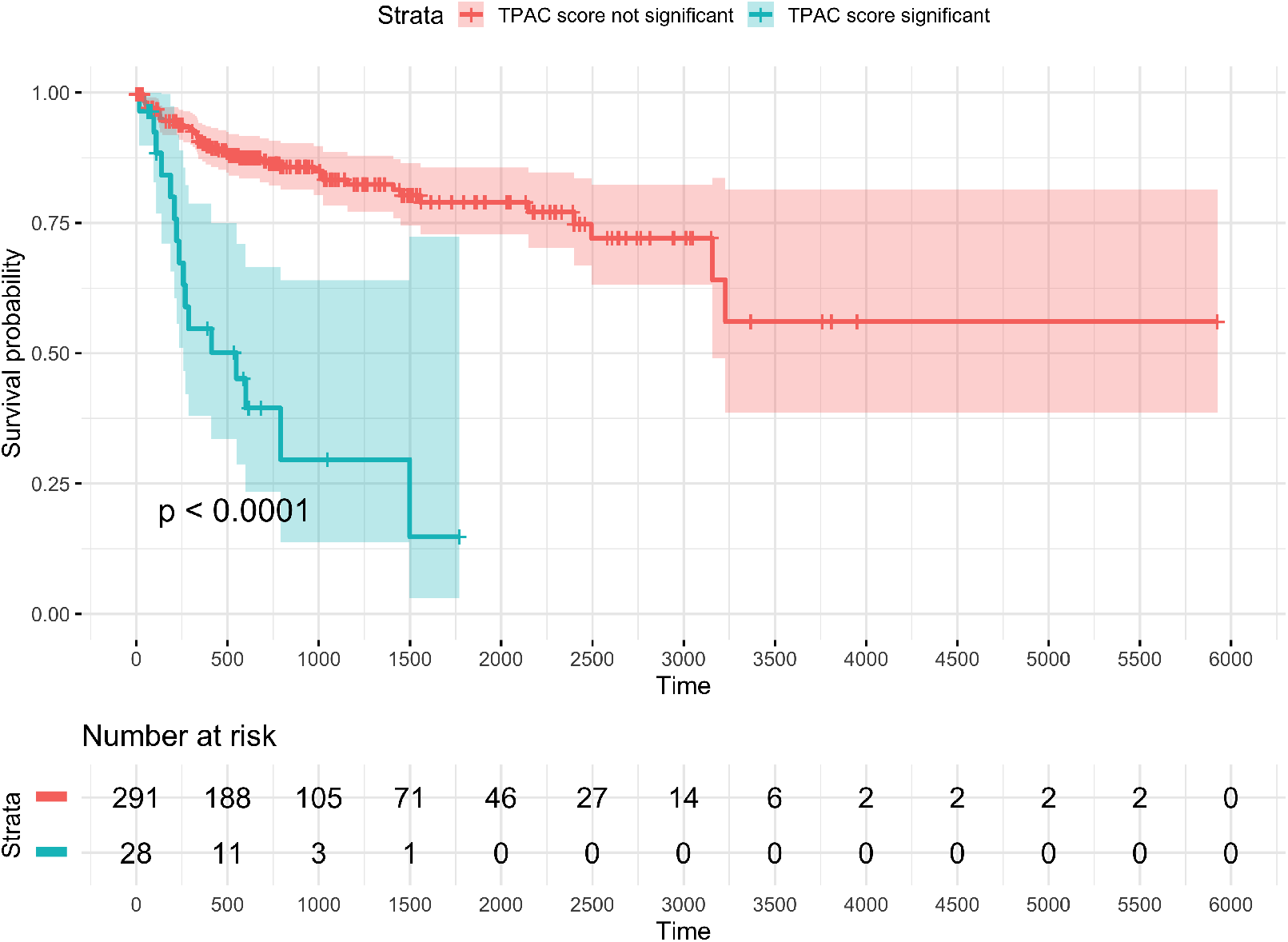
Kaplan-Meir plot for TCGA KIRP cohort and progression-free interval (PFI) outcome with patients stratified according to the significance of TPAC score for the MSigDB Hallmark MYC Targets V1 pathway. Significance was determined according to whether the FDR value associated with the TPAC score was *<* 0.25 where the family of hypotheses included the TPAC scores for all 50 Hallmark pathways for all 321 KIRP samples with PFI data (16,050 total hypotheses).

## 4 Conclusion

Cancer biology is highly tissue-specific: most cancer-driving somatic alterations occur in just a limited number of tissues and inherited mutations frequently have a tissue-specific functional impact. As we have explored in prior work [19], cancer tissue-specificity can be leverged to improve the power and accuracy of cancer genomic analyses. To leverage the associations between gene activity in normal and malignant tissue for pathway analysis, we have developed a new single sample pathway analysis method for tumor-derived transcriptomics data named TPAC (tissue-adjusted pathway analysis of cancer). The TPAC method uses the normal tissue-specificity of human genes to compute a robust multivariate distance score that quantifies pathway dysregulation in each profiled tumor.Because the null distribution of the TPAC scores has an accurate gamma approximation, both population and sample-level inference is supported. As demonstrated through an analysis of TCGA RNA-seq data, TPAC pathway scores are more strongly associated with both patient prognosis and tumor stage than the scores generated by existing single sample pathway analysis methods. The companion website (https://hrfrost.host.dartmouth.edu/TPAC) provides access an R package mplementation of the TPAC method that supports analysis of gene expression data for cancers associated with the 18 normal human tissues listed in Table 1.

